# Revisiting the role of beta-tubulin in *Drosophila* development: beta-tubulin60D is not an essential gene, and its novel *Pin*^*1*^ allele has a tissue-specific dominant-negative impact

**DOI:** 10.1101/2021.09.29.462296

**Authors:** Ramesh Kumar Krishnan, Naomi Halachmi, Raju Baskar, Bakhrat Anna, Adi Salzberg, Uri Abdu

## Abstract

Diversity in cytoskeleton organization and function may be achieved through alternative tubulin isotypes and by a variety of post-translational modifications. The *Drosophila* genome contains five different β-tubulin paralogs, which may play an isotype tissue-specific function *in vivo*. One of these genes, the *beta-tubulin60D* gene, which is expressed in a tissue-specific manner, was found to be essential for fly viability and fertility. To further understand the role of the beta-tubulin60D gene, we generated new *beta-tubulin60D* null alleles (*beta-tubulin60D*^*M*^) using the CRISPR/Cas9 system and found that the homozygous flies were viable and fertile. Moreover, using a combination of genetic complementation tests, rescue experiments, and cell biology analyses, we identified *Pin*^*1*^, an unknown dominant mutant with bristle developmental defects, as a dominant-negative allele of *beta-tubulin60D*. We also found a missense mutation in the *Pin*^*1*^ mutant that results in an amino acid replacement from the highly conserved glutamate at position 75 to lysine (E75K). Analyzing the β-tubulin structure suggests that this E75K alteration destabilizes the alpha-helix structure and may also alter the GTP-Mg^2+^ complex binding capabilities. Our results revisited the credence that *beta-tubulin60D* is required for fly viability and revealed for the first time in *Drosophila*, a novel dominant-negative function of missense *beta-tubulin60D* mutation in bristle morphogenesis.

**Author summary:** Diversity in cell microtubule cytoskeleton organization and function may be achieved through alternative tubulin isotypes and by a variety of post-translational modifications. The expression pattern of different tubulin isotypes (both α and β subunits) can vary according to cell type and stage of development, which contribute significantly to cell-specific MT organization and function. In this study, we revisited the role of one of the beta-tubulin isotopes in *Drosophila*, namely, beta-tubulin60D. This is the first study where a well molecularly defined protein null allele of *βTub60D* was generated and characterized. This well-characterized *βTub60D* allele demonstrated unambiguity that *βTub60D* is not an essential gene, as was described before. Moreover, we identified *Pin*^*1*^, an unknown dominant mutant with bristle developmental defects, as a dominant-negative allele of *beta-tubulin60D*. We also found a missense mutation in the *Pin*^*1*^ mutant that results in an amino acid (E75K). Analyzing the β-tubulin structure suggests that this E75K alteration destabilizes the alpha-helix structure and may also alter GTP-Mg^2+^ complex binding capabilities. Thus, our results also revealed for the first time in *Drosophila*, a novel dominant-negative function of a missense *beta-tubulin60D* mutation, which has a tissue-specific function.

## Introduction

Microtubules are polymers of α/β tubulin subunits, and they carry out a wide range of functions in eukaryotes (1)(2)(3). The expression of different tubulin isotypes can vary according to cell type and stage of development (4). The *Drosophila β-tubulin* gene family includes five members, each expressed in a unique pattern based on developmental timing and tissue-type specificity (5). The most divergent β-tubulin paralogs (*βTub85D* and *βTub65B*) are expressed exclusively in testis. *βTub85D*, but not *βTub65B*, has been characterized in considerable detail; it is required in the germline for male meiotic divisions and sperm axoneme formation (6,7). *βTub56D* is maternally supplied to the embryo and zygotically expressed during neurogenesis and in muscle attachment sites shortly after the insertion of muscles into the epidermis (7–10). It was shown that *βTub56D* is required for myoblast fusion, myotube elongation, and sarcomere formation during *Drosophila* embryogenesis (11). Recently, it was shown that the *βTub97EF* function is dispensable for viability and fertility, but it has a tissue-specific requirement for regulation of MT stability in a temperature-dependent manner. The expression of *βTub60D* is also tissue-specific; during embryogenesis, *βTub60D* expression starts in differentiating mesodermal cell types and occurs in chordotonal organs, imaginal discs, and somatic cells of the adult gonads (12,13). An extensive study on the role of *βTub60D* led to the identification of multiple alleles of *βTub60*, which showed lethality at different stages of development, from embryogenesis to larval stages (13–15).

Microtubules serve both as a scaffold for intracellular transport and contribute to cell polarity (16). The elongated *Drosophila* bristle is a single polyploid, highly polarized cell with a distinct direction of growth and a cone-like shape (17,18). The polarized *Drosophila* mechanosensory bristle cytoplasm is filled with short MTs that constitute a significant component of the shaft cytoplasm. These MTs appear to be stable during development and shorter in length than the mature bristle shaft (19). MT organization in bristles revealed two populations of MTs: one population is stable and uni-polarized, organized with their minus-end toward the bristle tip (20), and believed to serve as polarized tracks for proper organelle and protein distribution (19). The second MT population is dynamic with mixed polarity and contributes to proper axial growth (20,21), probably establishing bristle polarity (22).

This study reveals that *β-tubulin60D* is not an essential gene, as was described before, and elucidates a tissue-specific role of one of the *β-tubulin* paralogs, *β-tubulin60D*, in bristle MT assembly. We identified *Pin*^*1*^ as a dominant-negative allele of *β-tubulin60D*, which explicitly affects bristle development. Using sequencing and structural analysis, we demonstrated that the *Pin*^*1*^ mutation is caused by a single amino acid substitution, which affects the GTP-Mg^2+^ complex binding and interferes with the alpha helix’s stability.

## Results

To identify genes that may be involved in bristle development, we compared the repertoire of proteins of Su(H); sca-Gal4 flies, which lack all their bristles, versus wild-type flies (Fig. 1A). This comparative proteomic analysis/approach allowed us to identify bristle-specific proteins. We generated flies lacking bristles on their thorax by upregulating Su(H) expression specifically in the bristle lineage. Su(H) overexpression resulted in a complete loss of bristle cells (both microchaeta and macrochaeta) and the formation of extra socket cells (23) (Figure 1B). Proteins were extracted from thoraces of wild-type flies and Su(H) overexpressing, digested by trypsin, and analyzed by LC-MS/MS on Q Exactive Plus (Thermo). Samples were prepared/analyzed in triplicates for statistical significance. The complete list of differentially expressed proteins is presented in Supplementary File 1. Specifically, our proteomic analysis identified 27 proteins that were significantly downregulated in the flies expressing Su(H) as compared to wild type (Fig. 1C, D). Among these differentially expressed proteins, β-tubulin-60D showed the most significant (*p*-value<3.18E^-05^) fold change. We, therefore, decided to study the role of β-tubulin-60D in bristle development.

**Figure 1.**
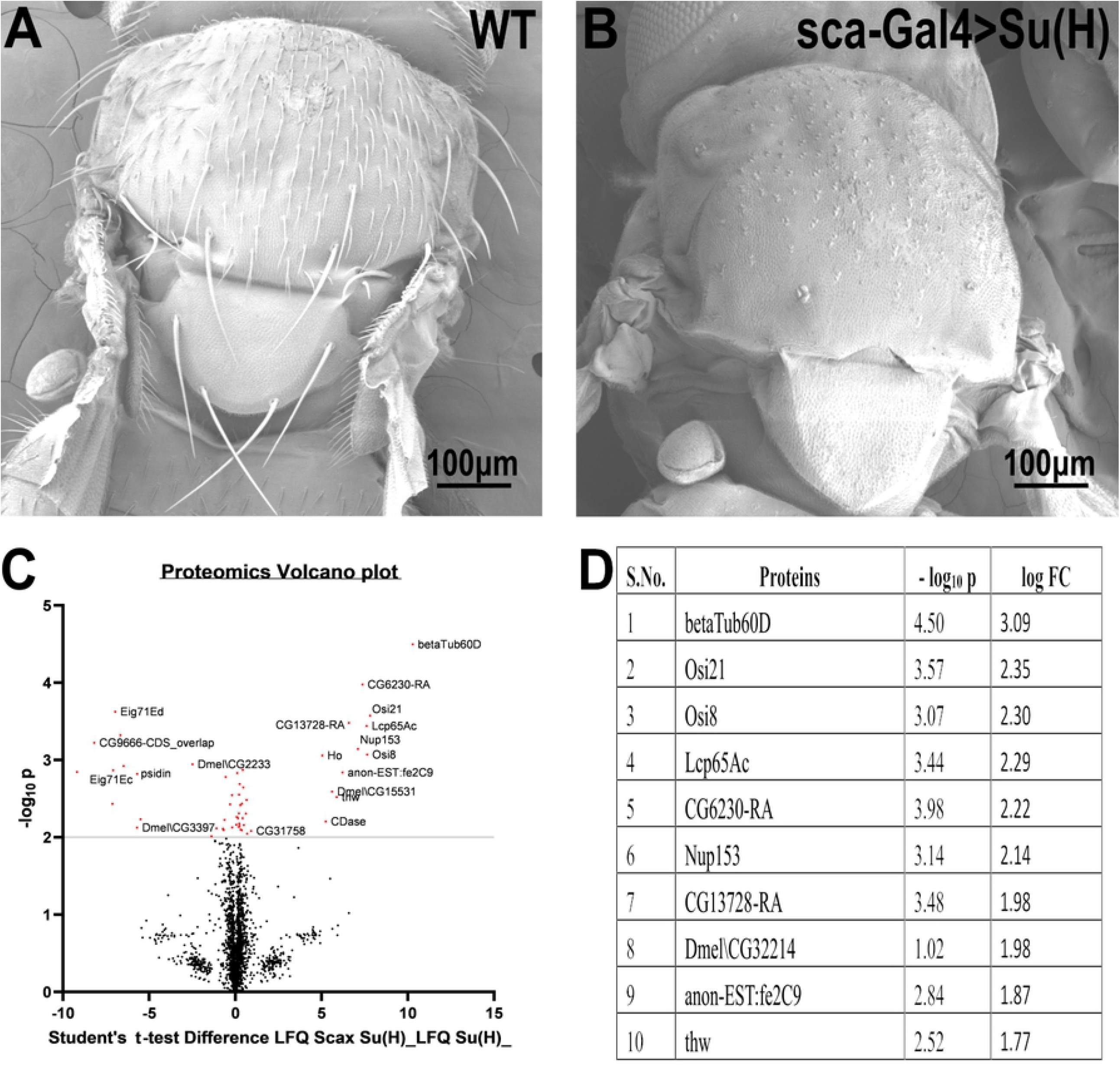
Proteomic profiling in the thoracic tissue of *Drosophila*. Scanning electron micrographs of the thorax wild-type (A) and sca-Gal4>Su (H) (B). Suppressor of Hairless [Su (H)], a transcriptional regulator in the notch signaling pathway, when driven by a sca-Gal4 driver, results in the complete absence of both microchaeta and macrochaeta. (C) Volcano plot showing differentially expressed proteins between WT (control) and Su (H); Sca Gal 4 (test) groups. Proteins with statistically significant differential expression (-log_10_ p > 2.0) are located in the top right and left quadrants. (D) Quantitative proteomics table of proteins with differential abundance in biological triplicates. Twenty-seven proteins were found to be downregulated as compared to the wild-type. The table shows the first ten proteins with their –log10 p-values and corresponding log-fold change values. *β-tubulin 60D* was chosen as a candidate protein because of its highly significant p-value.

### *β-tubulin60D* is expressed exclusively in the bristle shaft

To analyze the distribution/expression pattern of *β-tubulin60D* protein within the bristle lineage, we immunostained pupal thoraces using *β-tubulin60D*-specific antibodies. This staining revealed that *β-tubulin60D* protein is distributed along the entire bristle shaft but is excluded from the other lineage-related cells, namely, the socket, neuron, and sheath cells (Fig. 2A–C).

**Figure 2.**
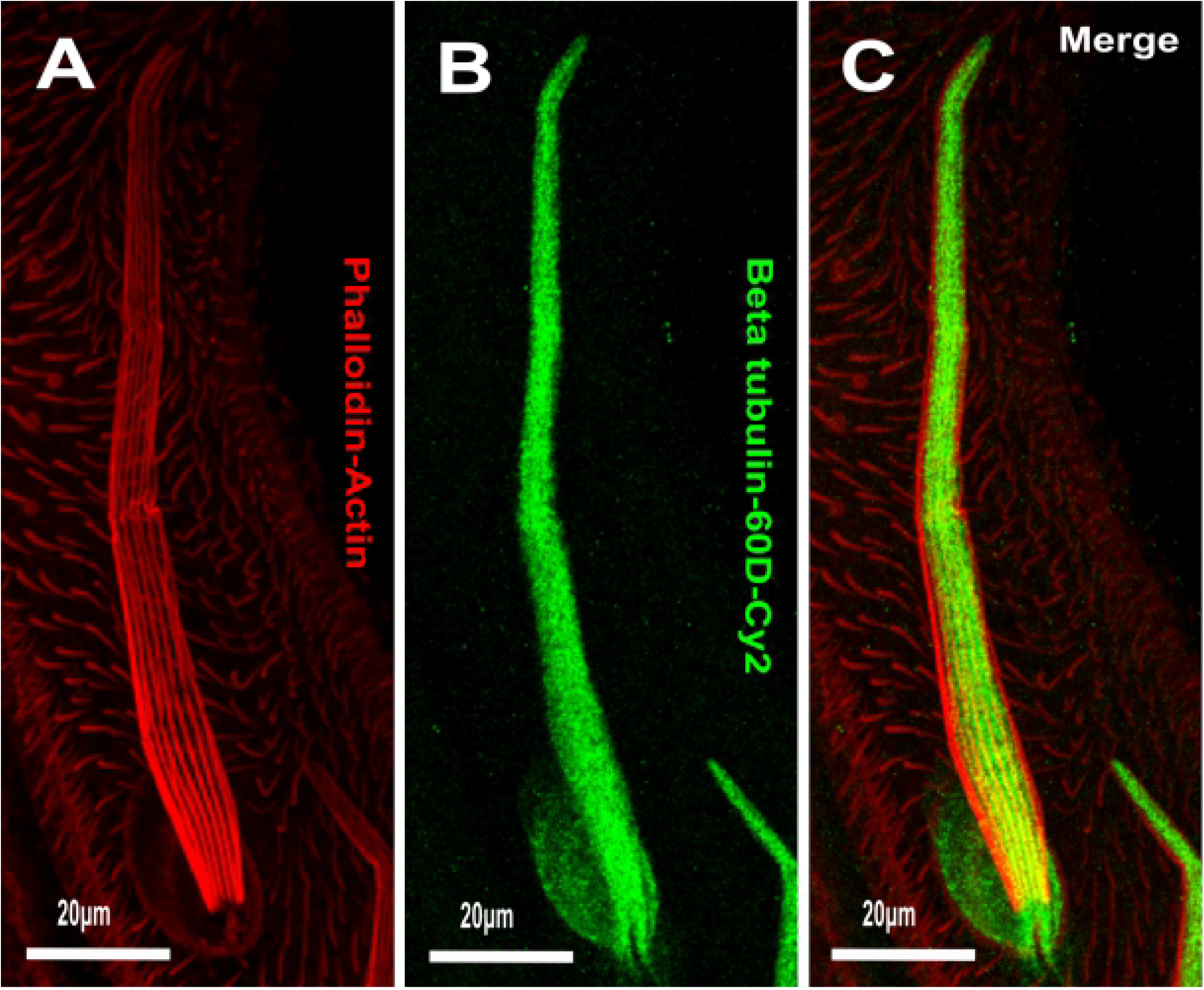
Expression of *β-tubulin-60D* in bristle. Confocal projections of bristles of ∼38 h APF from wild-type (A–C) pupae stained with Oregon red-phalloidin (red) and with *anti-β-tubulin-60D* antibodies (green). In wild-type bristles, *β-tubulin-60D* is abundant along the entire bristle shaft. APF – after prepupa formation.

### Loss of *β-tubulin60D* does not lead to lethality or sterility and does not impair bristle development and ChO morphogenesis

To test whether *β-tubulin60D* is involved in bristle development, we generated a *β-tubulin-60D* null allele using CRISPR/Cas9-mediated genome editing. For our CRISPR experiment, we designed two sgRNAs to replace the second exon by inserting the visible marker 3 X-P3-dsRed (Fig. 3A). Five independent mutated knockin insertion *Drosophila* lines were generated and named: *β-tubulin60D*^*M1*^ to *β-tubulin60D*^*M5*^. First, we confirmed that all the five mutated lines contained the 3 X-P3-dsRed, which replaced the entire second exon (Fig. 3B). To verify that indeed *β-tubulin60D*^*M*^ is a protein null allele, we stained homozygous larvae and pupae and examined *β-tubulin60D* expression in the chordotonal organ and bristle. As described above, usually, *β-tubulin60D* protein is expressed both in the bristle shaft (Fig. 4A–C) and in the cap cell of the ChO (Fig. 4G–I). However, in the *β-tubulin60D*^*M*^ mutant, no expression was detected in the bristle (Fig. 4D–F) or in the ChO cap cells (Fig. 4J–L), confirming that we had generated a complete loss of function allele of the *β-tubulin60D* gene. Previously, it was published that the *β-tubulin60D* gene is essential for viability and fertility (13). In contrast to this published data, we found that all our *β-tubulin60D*^*M*^ alleles were viable, and both males and females were fully fertile (Supplementary Table 1).

**Figure 3.**
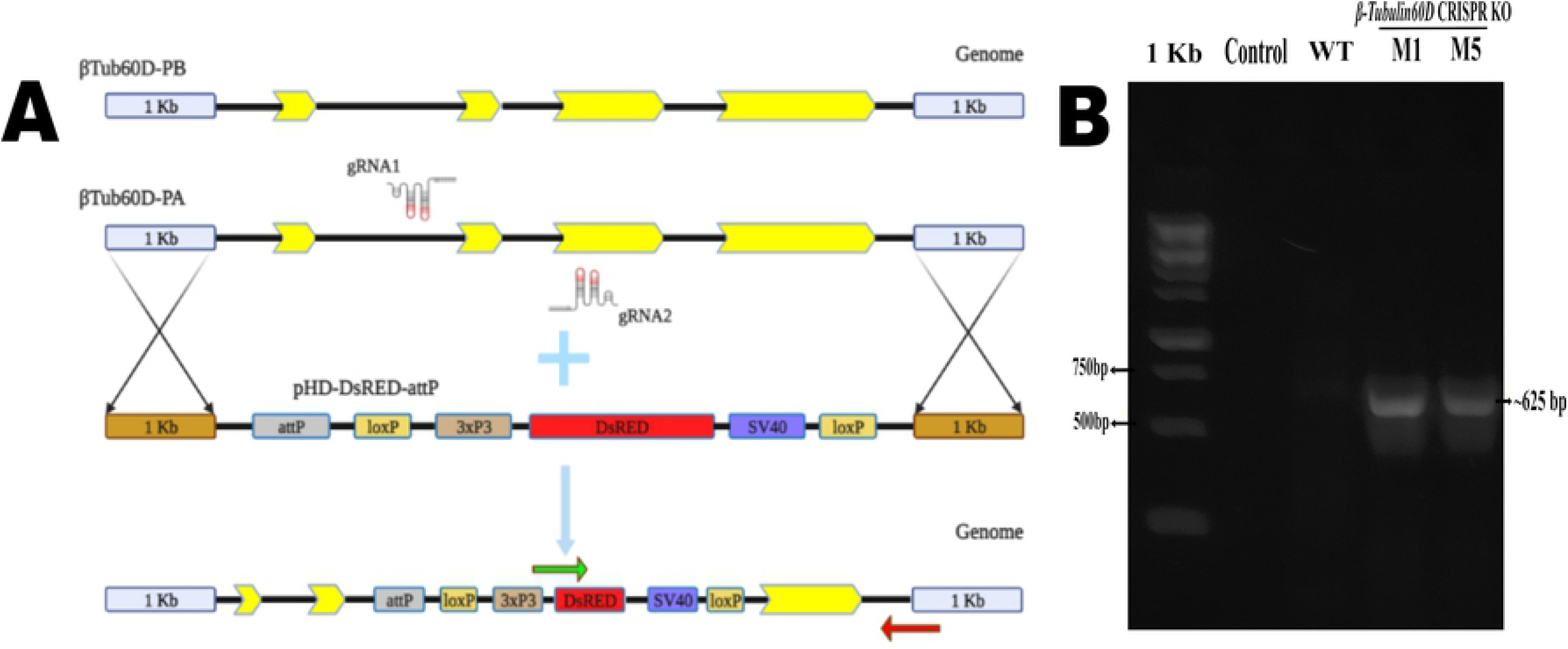
(A) Schematic diagram showing the nature of *beta-tubulin* loci and generation of β-tubulin KO (*β-tubulin60D*^*M*^) flies using CRISPR-mediated homology-directed repair (HDR) with the donor pHD-DsRed-attP vector. All two β-tubulin isoform exons are shown in yellow on loci, shown in black. Green and red arrows represent the binding sites of the forward and reverse primers used for genotyping, respectively. (B) PCR analysis of the genotyping of *β-tubulin60D*^*M*^ flies. The ∼625 bp band in the β-tubulin KO lane (*β-tubulin60D*^*M1*^ and *β-tubulin60D*^*M5*^) depicts the deletion of the second exon of β-tubulin from the genome and its replacement with DsRed using the primers depicted in (4A).

**Figure 4.**
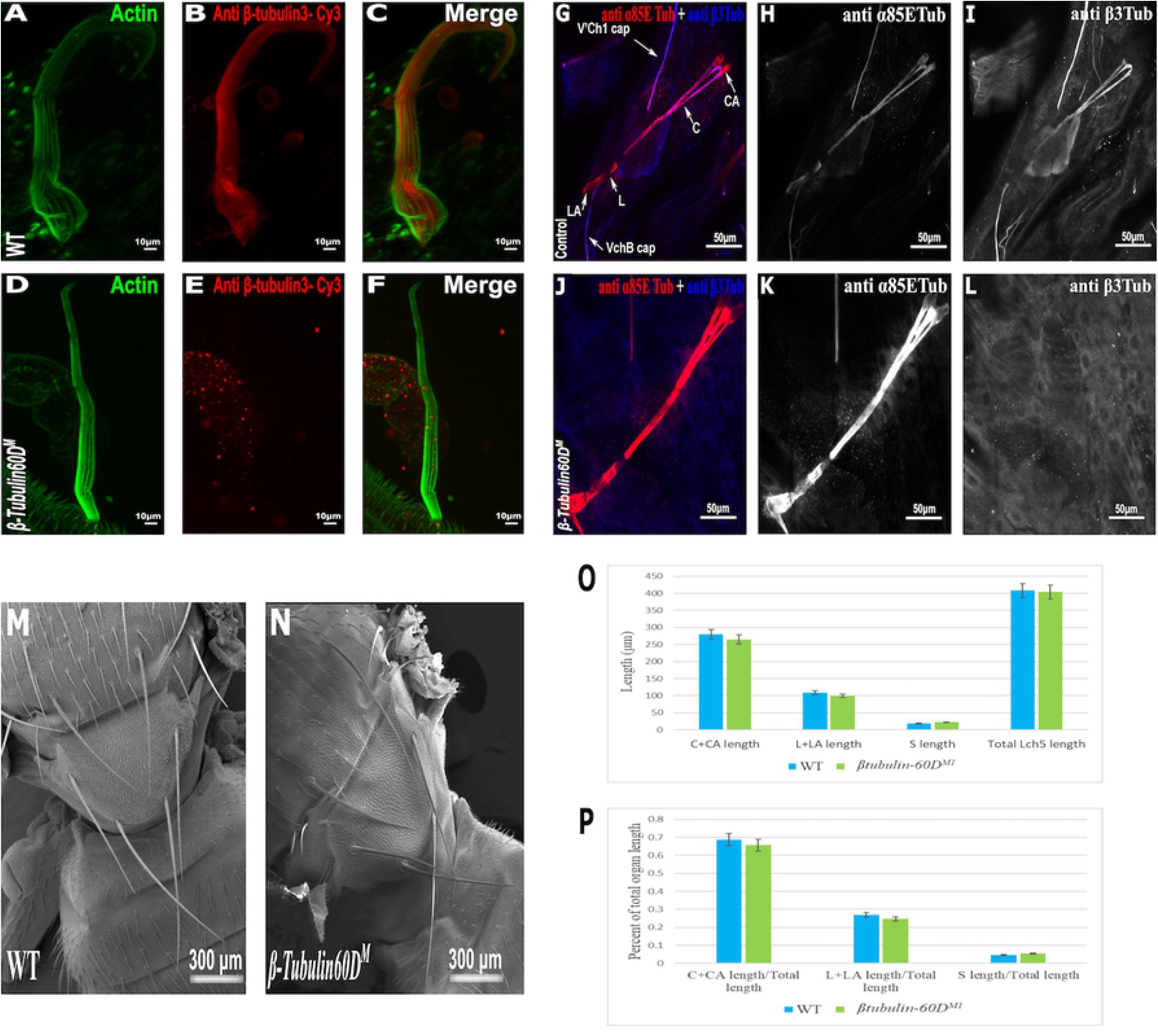
*β-tubulin-60D* null allele shows the absence of β3-tubulin in both bristles and chordotonal organs. Confocal projections of bristles of ∼38 h APF from wild-type (A–C) and *β-tubulin60D*^*M*^ (D–F) pupae stained with Oregon green-phalloidin (green) and with anti-β3-tubulin antibodies (red). In wild-type bristles, *β-tubulin-60D* is abundant along the entire bristle shaft, whereas in *β-tubulin60D*^*M*^ flies, there is a complete absence of the β3-tubulin. APF – after prepupa formation. Immunostaining of an LCh5 organ of a third instar WT (G–I) and β-tubulin-60D CRISPR KO (J–L) larvae by double staining with anti-α85ETub antibodies (in red) and *anti-β-tubulin60D* (in blue). *Anti-β-tubulin60D* is expressed explicitly in cap cells (I), whereas it is not present in the cap cells of *β-tubulin-60D* CRISPR KO lines (L). ***β-tubulin60D***^***M***^ **mutants show normal bristle development**. Scanning electron micrograph of adult bristles from wild-type flies (M) and *β-tubulin60D*^*M*^ (N). The CRISPR mutants do not show any visible structural defects in the bristle and appear similar to WT bristles. **β3-tubulin does not play a significant role in chordotonal organ morphogenesis**. The length of the different ChO cells of LCh5 organs of *β-tubulin60D*^*M1*^ homozygous third instar larvae was measured and compared to the wild-type larvae. (O) The graph shows the average length (in μm) of the cap (C) + cap-attachment cells (CA), ligament (L) + ligament-attachment cells (LA), and space (S) between the cap cells and ligament cells (this space corresponds to the scolopale cell). (P) The length of each cell was normalized to the total length of the organ. No significant difference is seen in both homozygous and heterozygous *β-Tubulin60D* compared to the wild-type larvae.

The specific expression of *β-tubulin60D* in the bristle shaft cells and its dramatic down-regulation protein in bristle-less flies could point to a possible role of this gene in bristle development. To address this issue, we examined bristle morphology in *β-tubulin60D*^*M*^ flies by scanning electron microscopy. Surprisingly, no apparent defects were detected in the bristles of the *β-tubulin60D*^*M*^ mutant/homozygous flies (Fig. 4 M, N).

Since *β-tubulin60D* is expressed solely in the ChO cap cell—the only cell type within the ChO lineage that elongates dramatically during larval growth—we investigated the possible role of *β-tubulin60D* in cap cell elongation by measuring the length of the different ChO cells in wild-type versus mutant third instar larvae. We found that, on average, the length of cap plus cap-attachment cells in wild-type larvae was 280 ± 38.9 mm (n=42), constituting 68.5% of the organ’s total length. In *β-tubulin60D*^*M1*^ homozygous larvae, no significant change in the length of the cap plus cap-attachment cells was noticed; 265.5 ± 39.9 mm (n=59), constituting 68.9% of the organ’s total length (Fig. 4 O–P). These results suggest that *β-tubulin60D* does not play a significant role in cap cell elongation and ChO morphogenesis.

### *Pin*^*1*^ is *a* dominant-negative allele of *β-tubulin60D*, which affects bristle but not ChO development

In parallel to the generation of the *β-tubulin60D* null allele, we searched known but uncharacterized mutations that cause abnormal bristle phenotypes for mutations that map to the genomic region in the vicinity of the *β-tubulin60D* gene (thus representing candidate alleles of *β-tubulin60D*). One such bristle defective mutant is called *Pin*. In the heterozygous *Pin*^*1*^ allele, macrochaeta, but not microchaeta, are shortened and sharply tapered at the tip (Fig. 5B, B’) compared to wild-type bristles (Fig. 5A, A’). The length of macrochaeta from heterozygous *Pin*^*1*^ mutant (Fig. 5B) measured 224.87±16.9 μm, which was significantly shorter (*p* < 0.01) as opposed to the wild-type, which had a bristle length of 395.79±1.9 μm (Table 1). Homozygous *Pin*^*1*^ mutants (Fig. 5C, C’) die as pharate adults, and the average length of their bristles is significantly shorter (40.73±4.2 μm) than the heterozygous *Pin*^*1*^ mutant.

**Table 1.** Table showing the length, base width, and tip width of wild-type along with the homozygous, heterozygous, hemizygous, and transheterozygous Pin^1^ mutants. Tukey’s test for post-hoc analysis shows that bristle length is statistically significant to each other (*p* < 0.01) compared to WT. When the transheterozygous mutant is rescued with a wild-type *β-tubulin-60D*, the bristle length equals almost that of the WT bristles. Tukey’s test for post-hoc analysis shows that bristle length is the same statistically, thereby showing that *β-tubulin-60D* rescues the bristle phenotype in the mutants. Ten SEM micrographs of the thoracic tissue were taken for analysis, and a minimum of five bristles was taken for measurement. ^a-d^ Different letters in the same column show that they are statistically significant compared to the wild-type, whereas the same letter in the column shows no significance.

| S.No. | Genotype | Length ($\mu\text{m}$) | Base width ($\mu\text{m}$) | Tip width ($\mu\text{m}$) |
| --- | --- | --- | --- | --- |
| 1 | WT | 395.79±1.9<sup>a</sup> | 9.90±0.9<sup>a</sup> | 3.87±0.2<sup>a</sup> |
| 2 | Pin<sup>1</sup>/Pin<sup>1</sup> | 40.73±4.2<sup>b</sup> | 6.99±0.4<sup>b</sup> | 1.90±0.2<sup>b</sup> |
| 3 | Pin<sup>1</sup>/CyO | 224.87±16.9<sup>c</sup> | 11.20±0.5<sup>a</sup> | 4.29±0.2<sup>a</sup> |
| 4 | Pin<sup>1</sup>/Df | 57.57±5.6<sup>d</sup> | 9.75±0.5<sup>a</sup> | 2.77±0.1<sup>a</sup> |
| 5 | Pin<sup>1</sup>; β-Tubulin60D<sup>M</sup> | 76.57±6.3<sup>f</sup> | 8.53±0.40<sup>a</sup> | 2.11±0.21<sup>c</sup> |
| 6 | Pin<sup>1</sup>/ β-Tubulin60D<sup>M</sup>; neur> β-tubulin-60D | 328.17±8.6<sup>a</sup> | 8.75±0.1<sup>a</sup> | 3.19±0.2<sup>a</sup> |

**Figure 5.**
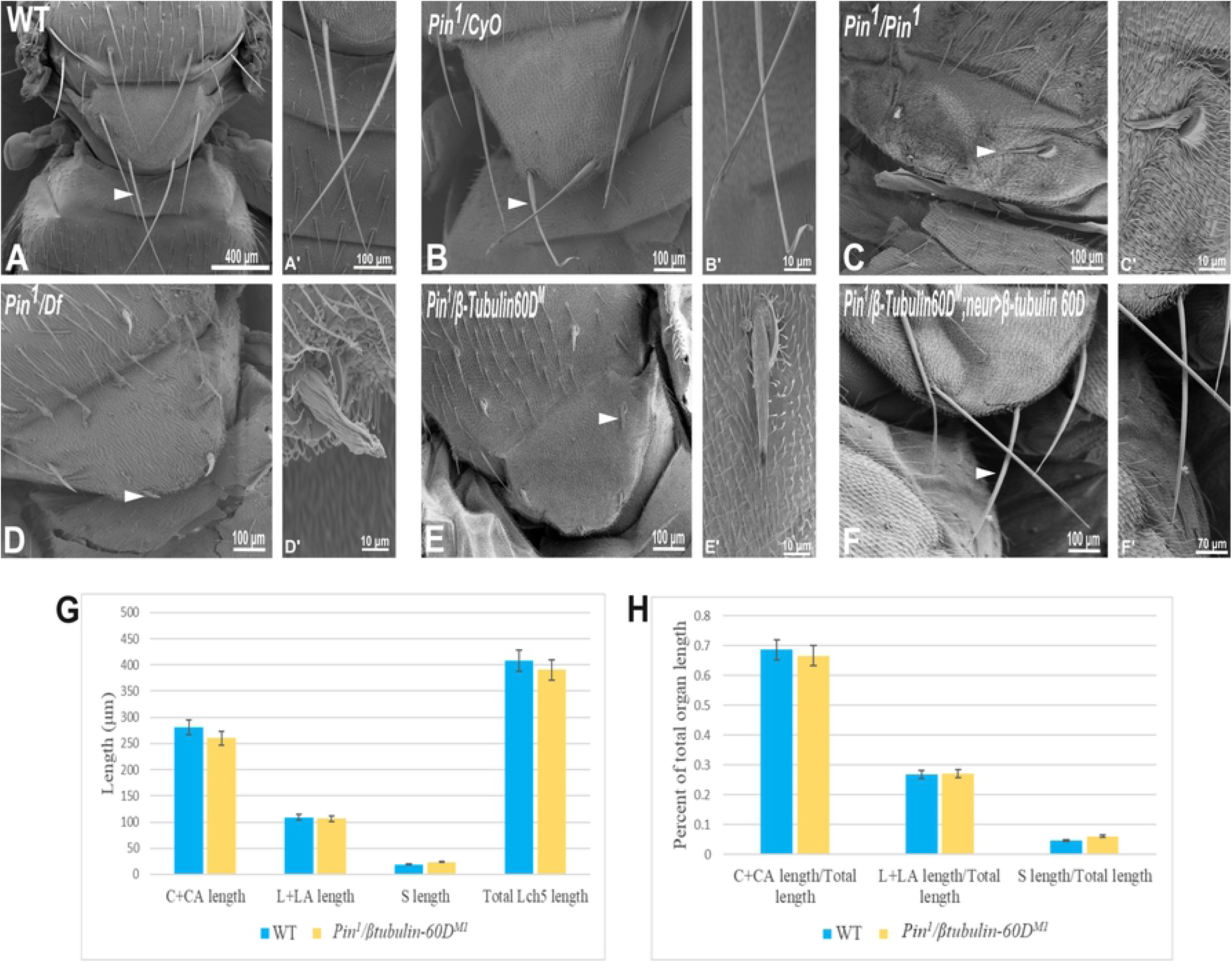
Gene mapping of Pin^1^ bristle phenotype. Scanning electron micrograph of adult bristles from wild-type flies (A), Pin^1^/CyO (B), Pin^1^/Pin^1^ (C), Pin^1^/Df (D), Pin^1^/*β-tubulin60D*^*M*^ (E), and Pin^1^/*β-tubulin60D*^*M*^; neur>β-tubulin60D. The Pin^1^ bristle (B’) is shorter compared to wild-type bristles (A’). Also, Pin^1^/Df bristle (D’) and Pin^1^/*β-tubulin60D*^*M*^ (E’) bristles are comparatively smaller than Pin^1^ and wild-type bristles. Pin^1^ and Pin^1^/Df have abnormally organized surface grooves. Pin^1^/*β-tubulin60D*^*M*^ bristles show smoother surfaces. ***β-tubulin-60D* rescues the bristle phenotype of *Pin***^***1***^. (F) shows the rescued bristle phenotype by *β-tubulin-60D*, resulting in longer bristles with properly tapered tips (F’) just like the wild-type bristles. Thus, the expression of *β-tubulin-60D* using neur-Gal4 rescues the Pin^1^ bristle phenotype. Arrowheads point to the bristle, which is shown as a higher magnification image. The length of the different ChO cells of LCh5 organs of *Pin*^*1*^*/β-tubulin60D*^*M1*^ transheterozygous third instar larvae was measured and compared to the wild-type larvae. (G) The graph shows the average length (in μm) of the cap (C) + cap-attachment cells (CA), ligament (L) + ligament-attachment cells (LA), and space (S) between the cap cells and ligament cells (this space corresponds to the scolopale cell). (H) The length of each cell was normalized to the total length of the organ. No significant difference is seen in both homozygous and heterozygous *β-tubulin60D* compared to the wild-type larvae.

To map the *Pin*^*1*^ allele, we first used deficiency mapping and found that *Df(2R)Exel6082*, which lacks the genomic region 60C4 to 60C7, fails to complement the *Pin*^*1*^ allele, as demonstrated by the effect on bristle development where the length of the hemizygous allele was 57.57±5.6 μm (Fig. 5D, D’), similar in their length to the *Pin*^*1*^ homozygous flies. Also, we found that the hemizygous flies, *Pin*^*1*^/Df, are viable, which means that the lethality of *Pin*^*1*^ homozygotes is probably due to other mutations in the background of the stock. Next, in order to find a smaller genomic region that will fail to complement the bristle phenotype of *Pin*^1^, we used two *nervy* alleles, *nervy* ^*PDFKG1*^ and *nervy* ^*PDFKG38*^, that remove the genomic region between the following P-elements: *KG(2)06386* and *KG(2)04837* (24). Since these deficiencies fail to complement the *Pin*^*1*^ allele bristle defects, it suggests that *Pin* could be an allele of one of 11 genes, among them *β-tubulin60D*. To test whether *Pin*^*1*^ is a dominant allele of *β-tubulin60D*, we crossed *Pin*^*1*^ with our *β-tubulin60D*^*M1*^ allele and found that trans-heterozygous flies had shorter bristles (76.57±6.3 μm) similar in their length to both hemi- and homozygous *Pin*^*1*^ mutants (Fig. 5E, E’). These results suggest that *Pin*^*1*^ is a dominant allele of the *β-tubulin60D* gene. To further characterize the nature of the *Pin*^*1*^ allele and to verify whether it is a dominant-negative or a neomorph allele of the *β-tubulin60D* gene, a rescue experiment was conducted. We generated transgenic flies that over-express the *β-tubulin60D* protein in the bristle using the *neur-Gal4* driver in a trans-heterozygous mutant background (*Pin*^*1*^/*β-tubulin60D*^*M1*^). The rescue experiment demonstrated that over-expression of *β-tubulin60D* completely rescued the short-bristle phenotype detected in *Pin*^*1*^/*β-tubulin60D*^*M1*^ flies (Table 1) (Fig. 5F, F’), suggesting that indeed *Pin*^*1*^ is a dominant-negative allele of *β-tubulin60D*.

Since the phenotypical analysis of the *Pin*^*1*^ allele implicates *β-tubulin60D* as required for bristle development, we tested whether this allele affects ChO morphogenesis. To test whether the *Pin*^*1*^ mutation also affects ChO development, we characterized the ChOs of third instar larvae of *Pin*^*1*^*/β-tubulin60D*^*M1*^ larvae and compared them to WT larvae. Theis analysis showed that in *Pin*^*1*^*/β-tubulin60D*^*M1*^ larvae, the length of the cap plus cap-attachment cells was 260.4 ± 29.4 μm (n=44), constituting 66.6% of the organ’s total length, which is not significantly different from WT larvae (Fig. 5G, H). This observation suggests that the *Pin*^*1*^ allele has a tissue-specific effect impairing only macrochaeta development.

To get better insight into the molecular nature of the *Pin*^*1*^ allele, we sequenced the *β-tubulin60D* coding region from the genomic DNA of *Pin*^*1*^*/Df* flies and compared the sequence to that of the WT strain/allele. The sequencing revealed a missense mutation in *Pin*^*1*^*/Df* at base pair 223 of its mRNA resulting in a glutamate-to-lysine replacement at position 75 (E75K). Alignment of the *Drosophila β-tubulin60D* protein across organisms from humans to yeast revealed that E75 is highly evolutionarily conserved (Fig. 6A). Moreover, the alignment of *Drosophila β-tubulin60D* protein with the other four *Drosophila* β-tubulin paralogs *(β-tubulin56D, β-tubulin65B, β-tubulin85D*, and *β-tubulin97EF)* also revealed higher conservation of this specific glutamate (Fig. 6B). The *Drosophila β-tubulin60D* protein contains the highly conserved N-terminal guanine nucleotide-binding region, intermediate domain (paclitaxel binding site), and C-terminal domains that constitute the binding surface for MAPs and molecular motors such as kinesins and dynein. By analyzing the β-tubulin structure (*Drosophila melanogaster*, Tubulin beta-1 chain, PDB: 6TIY), the newly identified *β-tubulin60D* mutation (E75K) lies at the guanine nucleotide-binding region. In this position, E75 acts as an alpha helix N-cap stabilizing residue via its hydrogen bond to the alpha-helix backbone (Fig. 6C). Additionally, E75 creates hydrogen bonds with two water molecules which are part of the Mg^2+^ hydration shell. This magnesium is critical for the GTP-Mg^2+^ complex binding at the tubulin nucleotide-binding site (Fig. 6D).

**Figure 6.**
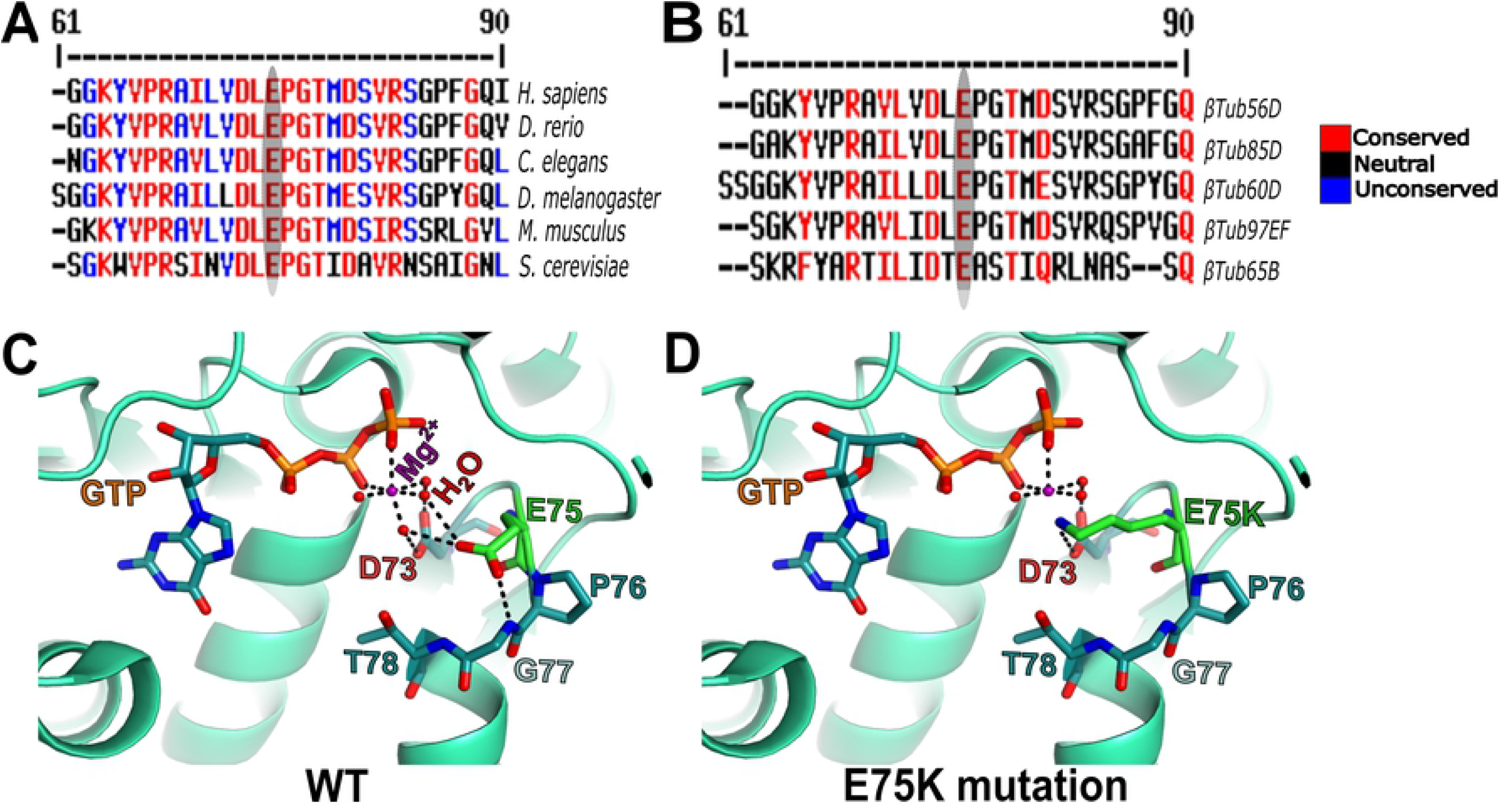
Amino acid conservation and structural modeling. Color scheme showing the sequence alignment of orthologues of *β-tubulin 60D* from different eukaryotes (A) and sequence alignment among all the known drosophila β-tubulin isoforms (B). The conservation scoring is performed by MultAlin. The scoring scheme works from 0 for the least conserved alignment position up to 10 for the most conserved alignment position, as indicated by the color assignments. The amino acid residue, glutamate {E}, is highly conserved among all the organisms ranging from human to *Drosophila*. The β-tubulin isoforms in *Drosophila* also show a similar degree of conservation. The conservation is highlighted with an ellipse. Structural comparison of wild-type *β-tubulin 60D* protein (C) and the Pin^1^ mutant *β-tubulin 60D* protein (D) shows how the single amino acid change at position 75 {E75K} affects the magnesium binding capacity of the protein, thereby affecting its functions.

### Microtubule network is mis-organized in *Pin*^*1*^ mutant

To further examine the effects of *Pin*^*1*^ on bristle shaft development, we characterized the organization of MTs in the bristle using antibodies against *β-tubulin60D* (25) and against acetylated tubulin (Fig. 7G–L), which recognize stable MT network in the bristle (22). Anti-*β-tubulin60D* staining revealed that the *β-tubulin60D* protein is present in *Pin*^*1*^ hemizygous pupae (Fig. 7D–F). This observation suggests that the E75K alteration found in *Pin*^*1*^ does not significantly affect the stability of the *β-tubulin60D* protein; still, this staining showed that MTs are extremely disorganized (Compare Fig. 7A–C to Fig. 7D–F). Whereas in WT pupae, MTs are found throughout the bristle shaft, in hemizygous *Pin*^*1*^ mutants, they are not evenly distributed and often appear as aggregates found at various locations along the bristle shaft. Disorganization of the stable α-tubulin MT network was also evident in the hemizygous *Pin*^*1*^ mutants] (Compare Fig. 7G–I to Fig. 7J–L).

**Figure 7.**
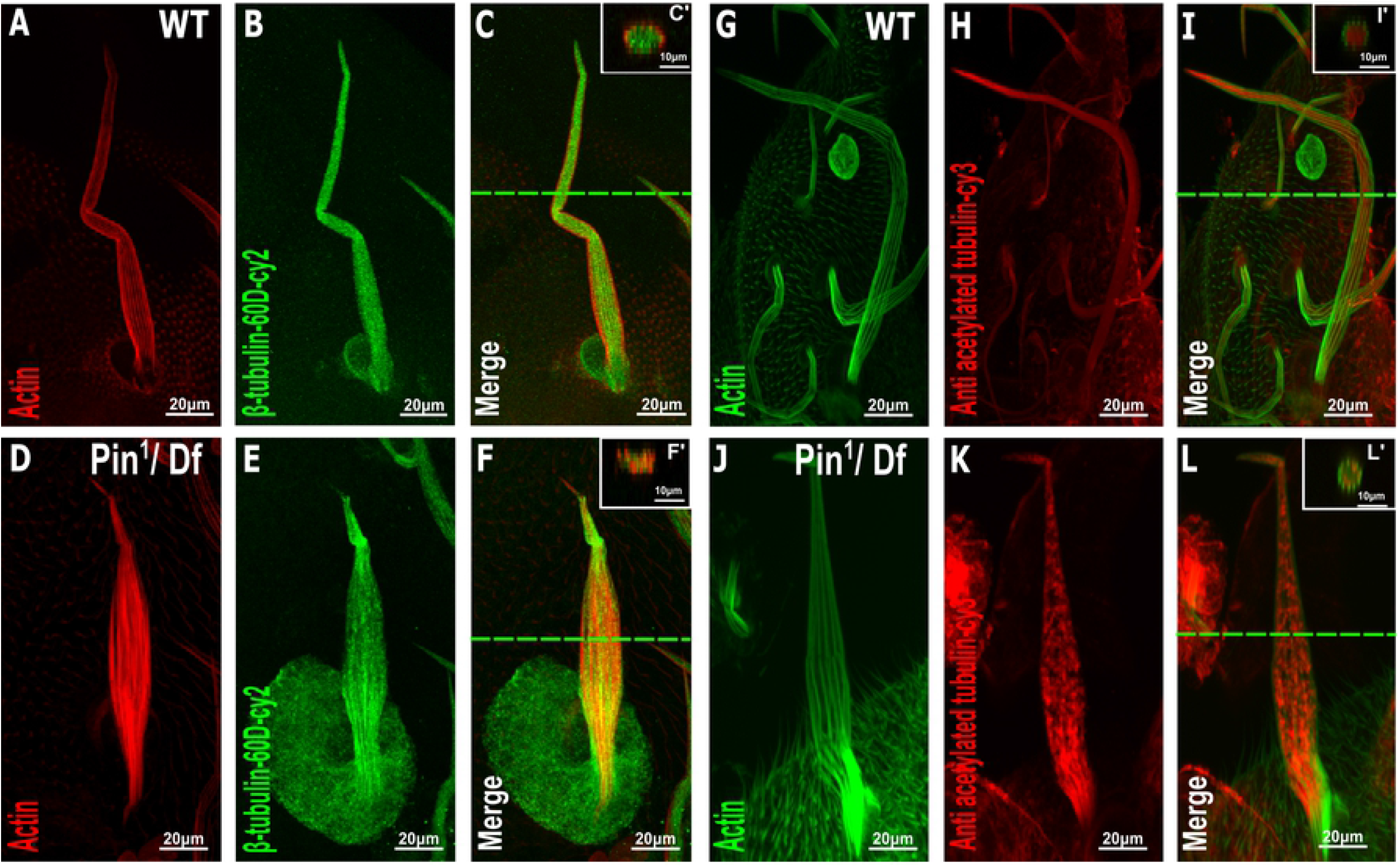
Distribution of β-tubulin and acetylated α-tubulin is affected in the Pin^1^ mutant bristle. Confocal projections of bristles of ∼37 h APF from WT (A–C) and Pin^1^/Df (D–F) pupae stained with red-phalloidin (red) and with *anti-β-tubulin-60D* antibodies (green). Digital cross-sections marked by a green line of wild-type (C’) and Pin^1^/Df (F’) pupae demonstrate a gradual decrease in *β-tubulin-60D* density at the middle of the bristle shaft. Confocal projections of bristles of ∼38 h APF from WT (G–I) and Pin^1^/Df (J–L) pupae stained with green-phalloidin (green) and with anti-acetylated tubulin-antibodies (red). Digital cross-sections marked by a green line of wild-type (I’) and Pin^1^/Df (L’) pupae demonstrate a patchy and uneven distribution in **a**cetylated α-tubulin density throughout the bristle shaft compared to that of the wild-type. APF – after prepupa formation. APF – after prepupa formation.

## Discussion

### *β-tubulin60D* is not an essential gene

This is the first study where a well molecularly defined protein null allele of *βTub60D* was generated and characterized. This well-characterized *βTub60D* allele demonstrated unambiguity that *βTub60D* is not an essential gene. These results disagree with previous studies in which multiple alleles of *βTub60* were generated, which showed lethality at different stages of development, from embryogenesis to larval stages (13–15). Using an ethyl methanesulfonate (EMS) or diepoxybutane (DEB) mutagen screen led to the identification of one larval lethal complementation group of five alleles, *β*3t^1^–*β*3t^5^, and some, but not all, of these alleles, could be rescued by a *βTub60* transgene. Examination of the homozygous and transhetrozygous phenotype suggested that *βTub60* is required for viability and fertility (13). In the second screen, eight new alleles of *βTub60* were identified; six were induced by EMS, one by gamma radiation, and one by P-element mutagenesis. All alleles were recessive lethal in the larval stages, except for two semi-lethal but sterile alleles (14). Some of the combinations of the transheterozygous alleles also exhibited bristle and flight defects. All of the alleles of *βTub60* that were generated by EMS showed that these alleles were not protein null (14). Sequencing one of the alleles, *β*3t2, which belongs to the class I severe alleles, revealed no lesion in the coding region of the gene (15). To this end, the *β*3t2 allele is the only available allele, but we found that it is no longer recessive lethal, and it complements *Pin*^*1*^ allele bristle defects (data not shown), suggesting that this stock is no longer a *βTub60* allele. To conclude, the facts that all other lethal alleles are not well molecularly characterized and also not available for further characterization, together with the fact that our molecularly defined protein null allele of *βTub60D* revealed that *βTub60D* is not an essential gene, led us to conclude that *βTub60D* is not required for *Drosophila* viability.

### *Pin*^*1*^ encodes a novel dominant-negative allele of the β-tubulin60D gene

To further characterize the *βTub60D* gene, we found that *Pin*^*1*^, an uncharacterized dominant bristle defects allele, is a novel dominant allele of the *β-tubulin60D* gene. First, our protein-null *βTub60D* alleles fail to complement the bristle defect found in hemizygous *Pin*^*1*^ mutants. Second, the expression of *βTub60D* protein specifically in the bristle completely rescues the bristle defect found in hemizygous *Pin*^*1*^ mutants. Third, as expected from the gene part of the MT lattice, the MT network in bristles from transhetrozygous and hemizygous *Pin*^*1*^ mutants is severely affected. Fourth, our genetic analysis showed a missense mutation in *Pin*^*1*^*/Df* at base pair 223 of its mRNA, resulting in an amino acid replacement from glutamate at position 75 to lysine (E75K). Bioinformatic analysis suggests that replacing the glutamate with lysine residue destabilizes the alpha helix since lysine is an alpha helix N-cap destabilizing residue. In addition, lysine’s positive charge will be located near the Mg^2+,^ which might prevent its binding or, in general, alter the GTP-Mg^2+^ complex binding capabilities.

In humans, mutations in β-tubulin genes are associated with defects in neuronal development (26–32), oocyte meiosis (33,34), thrombocytopenia (35), and macrothrombocytopenia (36,37). All these human mutations are found as heterozygous missense mutations, suggesting that either haploinsufficient or dominant-negative are the mechanism that causes these diseases. Study on the disease-associated mutations in *TUBB3* showed that R62Q, A302T, R380C, or R262C mutations impair tubulin heterodimer formation in vitro. The R62Q, R262H, R262C, A302T, and E410K mutations also disrupted microtubule dynamics in yeast. The E410, D417, and R262 mutations affect Kinesin binding to MT (29). These results suggest that these missense mutations do not affect *TUBB3* protein stability, but affect the cellular function of the MT network, maybe due to the “toxic” effect of the mutant tubulin isotype. However, the debate on the potential mechanisms for the disease-causing heterozygous tubulin mutants is still open. Our study showed that a complete loss of function of *βTub60D* does not affect fly viability, with no other obvious defects. The fact that the *βTub60D* null allele had no defects in *Drosophila* development, although it has tissue-specific expression, suggests that other *βTub* paralogues may compensate for the loss of the *βTub60D* gene. On the other hand, we demonstrated that *Pin*^*1*^ is a heterozygous missense allele of the *βTub60D* gene with a tissue-specific requirement. Thus, the fact that complete loss of o *βTub60D* had no apparent defects in *Drosophila* development and that *Pin*^*1*^ is a heterozygous missense allele of the *βTub60D* supports the idea that dominant-negative, but not haploinsufficient, is the mechanism underlying the function of the *Pin*^*1*^ allele.

## Materials and methods

### *Drosophila* stocks

Oregon-R was used as the wild-type control. The following mutant and transgenic flies were used: *Suppressor of Hairless, Su(H)* (38), *Df(2R)Exel6082* (Bloomington *Drosophila* Stock Center #7561), *nervy* ^*PDFKG38*^ and *nervy* ^*PDFKG1*^ (24), and *Pin*^*1*^ (Bloomington *Drosophila* Stock Center). For the rescue experiment, *M{UAS-βTub60D*.*ORF}ZH-86Fb* was used (39). Bristle-specific expression was induced under the control of the *neu-Gal4* driver (for the rescue experiment) or *Sca-Gal4* driver. All of the Gal4 lines were obtained from the Bloomington *Drosophila* Stock Center.

### Developmental staging and pupal dissection

Stages of all flies were determined from puparium formation (40). White prepupae were collected and placed on double-sided scotch tape in a petri dish placed in a 25°C incubator, as previously described (19). At the appropriate time of incubation (36 to 44 h APF, unless indicated otherwise), the pupae were dissected for live imaging, fixation, and proteomic screening. The pupal case was removed as described in (41). After removing the pupal case, the pupae were dissected as described elsewhere in detail (19).

### Bristle phalloidin and antibody staining

Bristle fixation and staining were performed as previously described (42,43). Confocal images were taken using an Olympus FV1000 laser scanning confocal microscope and are shown here as *z*-projections in a few optical frames that covered the bristle cell. Primary antibodies used were anti-α-acetylated tubulin mouse monoclonal antibodies (1:250; Sigma, T7451) and anti-β-tubulin mouse monoclonal antibodies (1:250; Sigma). Bristles from CRISPR KO flies were stained with anti-β-rabbit anti-β3Tub rabbit (1:1,000) (25). Cy3-conjugated goat anti-mouse (1:100; Jackson ImmunoResearch) secondary antibody was used. For actin staining, Oregon Green 488- or Alexa Fluor 568-conjugated phalloidin (1:250; Molecular Probes) was used.

### Scanning Electron Microscopy

Adult *Drosophila* flies were fixed and dehydrated by immersing them in increasing concentrations of ethanol (25%, 50%, 75%, and twice in 100%; 10 min each). The flies were then completely dehydrated using increasing concentrations of hexamethyldisilazane (HMDS) in ethanol (50%, 75%, and twice in 100%; 2h each). The samples were air-dried overnight, placed on stubs, and coated with gold. The specimens were examined with a scanning electron microscope (SEM; JEOL model JSM-5610LV). Length measurements of adult bristles were performed using Image J (version 1.52t) software with the straight-line tool. This tool allows the creation of line selections and then the calculation of the length of these lines. To test for differences in bristle length and width between the wild-type and the different mutants, we used a one-way analysis of variance (ANOVA) followed by a Tukey analysis.

### Sample preparation for mass spectrometry analysis

The pupal case was removed as described in (41). Then the pupae were dissected as described elsewhere in detail (19). The dissection procedure resulted in the isolation of thorax dorsal side tissue, which was then cleaned of interior organs and fat particles as described in (19). All procedures were conducted in phosphate-buffered saline (PBS). The head and abdomen parts of the tissue were cut, leaving only the thorax intact, which was then put in a vial of PBS with a protease inhibitor cocktail (Sigma). Each group consisted of triplicates of 20 thoracic tissues.

### Proteolysis and mass spectrometry analysis

The samples were ground in 10mM DTT 100mM Tris and 5% SDS, sonicated, and boiled at 95^0^C for 5 min. They were then precipitated in 80% acetone. The protein pellets were dissolved in 9M Urea and 100mM ammonium bicarbonate and reduced with 3mM DTT (60ºC for 30 min), modified with 10mM iodoacetamide in 100mM ammonium bicarbonate (room temperature for 30 min in the dark), and digested in 2M Urea25mM ammonium bicarbonate with modified trypsin (Promega), overnight at 37°C in a 1:50 (M/M) enzyme-to-substrate ratio. The resulting tryptic peptides were desalted using C18 tips (Harvard), dried, and re-suspended in 0.1% formic acid. They were analyzed by LC-MS/MS using a Q Exactive Plus mass spectrometer (Thermo) fitted with a capillary HPLC (easy nLC 1000, Thermo). The peptides were loaded onto a homemade capillary column (20 cm, 75 micron ID) packed with Reprosil C18-Aqua (Dr. Maisch GmbH, Germany) in solvent A (0.1% formic acid in water). The peptide mixture was resolved with a (5–28%) linear gradient of solvent B (95% acetonitrile with 0.1% formic acid) for 180 min, followed by a 15-min gradient of 28–95% and 15 min at 95% acetonitrile with 0.1% formic acid in water at flow rates of 0.15 μl/min. Mass spectrometry was performed in a positive mode using repetitively full MS scanning followed by high collision-induced dissociation (HCD, at 25 normalized collision energy) of the ten most dominant ions (>1 charges) selected from the first MS scan. The mass spectrometry data were analyzed using the MaxQuant software 1.5.2.8. (www.maxquant.org) using the Andromeda search engine, searching against the *Drosophila* UniProt database with a mass tolerance of 6 ppm for the precursor masses and six ppm for the fragment ions. Peptide- and protein-level false discovery rates (FDRs) were filtered to 1% using the target-decoy strategy. Protein tables were filtered to eliminate the identifications from the reverse database and common contaminants and single peptide identifications. The data were quantified by label-free analysis using the same software, based on extracted ion currents (XICs) of peptides, enabling quantitation from each LC/MS run for each peptide identified in the experiments. Statistical analysis of the identification and quantization results was done using Perseus 1.6.7.0 software

### Generation of *β-tubulin 60D* knockout flies by CRISPR Cas-9-mediated genome editing

To generate knockout flies, two guide RNA sequences were identified (sgRNA1 - GGCGGTCCCGTCTCCAAAGGGGG & sgRNA2 - GGAGCCCGGAACCATGGAGTCGG) at http://targetfinder.flycrispr.neuro.brown.edu/ and cloned into plasmid pU6-BbsI-chiRNA. Then 1 Kb sequence stretches upstream and downstream of *β-tubulin60D* were cloned into the donor pHD-DsRed-attP vector. Finally, injection of both vectors and fly screening was carried out by BestGene.

### Fertility assay

Three virgin *β-tubulin60D* ^*M*^ females were mated with two wild-types (WT) males, and two *β-tubulin60D* ^*M*^ males were mated with three virgin wild-type (WT) females in a vial containing yeast for two days. Matings were performed in triplicate for each genotype. The flies were transferred to new vials containing fresh yeast and were let to lay eggs for one day. The flies were then discarded, and the adult progeny were collected and counted after ten days at 25°C. The progeny per female and the average number and standard deviation of progeny per genotype were calculated from each vial. Finally, a percentage of relative fertility was calculated (44).

### Chordotonal organ staining/cuticle preparation

Seven larvae from each genotype were dissected and immune-stained as previously described (12). Mouse anti-αtubulin-85E (1:20) (45) was used for visualizing the LCh5 accessory cells and rabbit anti-βTub60D (1:1000) (25) for verifying the loss of βTub60D in the mutant larvae. Secondary antibodies were Cy3-conjugated anti-rabbit and Cy5-conjugated anti-mouse (1:100, Jackson Laboratories, Bar-Harbor, Maine, USA). Stained larvae were mounted in DAKO mounting medium (DAKO Cytomation, Denmark) and viewed using confocal microscopy (Axioskop and LSM 510, Zeiss). To guarantee similar age of all tested larvae, flies of all genotypes were let to lay eggs for 3 h, and the progeny were aged for 118–121 h at 24°C. For each chordotonal organ, we measured the length of the cap plus cap-attachment cells, the ligament plus ligament-attachment cells, and the space between the cap and ligament cells, which represents the scolopale cells. The length of each cell was normalized to the total length of the organ.

## Data Analyses

Quantitative data are expressed as mean ± standard error of the mean (SEM). All the statistical analyses were performed using a one-way ANOVA, and *p-*values ≤ 0.05 were considered significant for all analyses. Statistical significance was checked with a pairwise post-hoc Tukey HSD. All the statistical analyses were performed using STATISTICA, version 10.

## Acknowledgments

We thank Bloomington Stock Center, Christian Lehner, and Anette Preiss for generously providing fly strains. The research was supported by the Israel Science Foundation (ISF) (grant 278/16 to U.A and No. 674/17 to A.S).

## Supplementary data

**Supplementary File 1**. Summary of all the proteins identified in the proteomics screen.

**Supplementary Table 1.** β-tubulin-60D null allele does not affect male and female fertility. Tukey’s test for post-hoc analysis shows that the percentage of relative fertility has no significant difference when compared statistically to the control group. ^a^ Same letter in the column indicates no significant statistical difference between the groups.

| S.no. | Genotype | Average no. of progeny from 24 hr egg collection | % relative fertility |
| --- | --- | --- | --- |
| 1 | WT ♂ x WT ♀ | 98.33±0.7 | 100 <sup>a</sup> |
| 2 | CRISPR KO *β-Tubulin60D<sup>M</sup>* ♂ x WT ♀ | 94.66±1.1 | 96.2 <sup>a</sup> |
| 3 | CRISPR KO *β-Tubulin60D<sup>M</sup>* ♀ x WT ♂ | 87.66±1.3 | 89.1 <sup>a</sup> |

